# Development of membrane-like pre-stem cells after released from tube-shaped niches

**DOI:** 10.1101/2023.03.04.531096

**Authors:** Wuyi Kong, XiuJuan Han, XiaoPing Zhu, Hong Wang

**Affiliations:** Beijing Khasar Medical Technology Co., Beijing, China

**Keywords:** Lineage restriction, Bamboo-like inclusions, Membrane-like progenitors, Tube-shaped structures, Stem cell progenitors, DNA fragments

## Abstract

We previously identified three distinct pre-stem cell lineages that further develop into blood CD34-positive stem cells, and mesenchymal stem cells. The pre-CD34-positive stem cells are spore-like, while the pre-mesenchymal stem cells are fusiform-shaped. All of them originate or released from tube-shaped tissue structures, or niches. In the current study, we present two membrane-like pre-stem progenitors. One of them had red thin membrane structures that were from disconnection of bamboo-like tissues and developed into thin membrane-like multi-nucleated cells. The other membrane-like structure had geometric-shape and was the place of developing numerous c-kit-positive stem cell progenitors. One geometric-shaped membrane could produce dozens to hundreds of progenitors in a synchronized pattern. Our findings provide more evidence that postnatal stem cells are unipotent and self-renewed in sealed tube-shaped structures. The tube-shaped structures released their inclusions that have distinct morphological differences among the cellular lineages. Thus, the self-renewal of each lineage-distinct stem cell has its unique pattern. Further, our data suggests that postnatal stem cells are renewed via a recurrent pattern. Our findings again challenged the old concepts of stem cell niche components, the origins of stem cell lineages and the self-renewal of stem cells.

## Introduction

Stem cells are constantly needed in the body to replace aged cells and repair wounds. Thus, elucidating how and where stem cells are renewed has become a very important research topic. There seems to be a basic consensus that stem cells are either located or renewed in a microenvironment called a niche. Until now, most reports have described stem cell niches located in the bone marrow sinusoids, perivascular tissue or specific organs ^1^. They are composed of different types of cells ^2^ or composed of cells and extracellular structures such as the extracellular matrix (ECM) ^3,4^. However, these descriptions are not supported by any true images. In addition, the complexities and structures of these niches have remained largely unknown.

Most believe that stem cells in niches are in a quiescent state that can be activated by various factors or hormones and undergo self-renewal to maintain life-long production of mature cells ^5,6^. Reports on quiescent stem cells have been contradictory. Some reports indicated that the mechanism of postnatal stem-cell self-renewal is asymmetric division 7,8, and other reports based on imaging techniques in live mice indicated that dermal stem cells do not undergo asymmetric cell division during differentiation. Epidermal stem cells have equal potential to divide or directly differentiate ^9^. In addition, most reports regarding the function of stem cell self-renewal are based on counting cell colonies or the quantifying cell replication ability, which gradually leads stem cell self-renewal studies to cancer cells ^10,11^. Thus, the mechanism of postnatal stem cell self-renewal is still unclear, due to reported mechanisms are extremely varied.

The progenitor population has been described as downstream of haematopoietic stem cells (HSCs) that segregate blood cell lineages step by step, ultimately generating unipotent progenitor cells ^12-16^. However, a recent report claimed that lineage-restricted segregation occurs during early embryonic development ^17^, indicating that the cell lineages have been carefully categorized prenatally or that stem cells are switched from multipotent to unipotent during early embryonic development. Some reports indicate that HSCs are able to differentiate into restricted progenitors before cell division or that HSC transition to restricted blood progenitors occurs via transdifferentiation procedures ^18^.

Until now, the complete picture of the production of restricted blood progenitors is still unclear.

In our previous reports, we found that pre-stem tissues originate form tube-shaped structures. These pre-stem tissues have distinct morphological differences ^19^, such as thin filaments-released spore-like, or dark fragments-released fusiform-shape ^22^. Because each of these structures produces and releases only one distinct spore-like cell type, or progenitors, we believe that the lineage of these progenitors has been predetermined in their tube-shaped structure, or niche.

Here, we show two more morphologically distinct stem cell self-renewal procedures. Thin red membranes disconnected from red bamboo-like tissues and further develop into nucleated cells; and c-kit-positive small cell progenitors that were segregated from geometrically shaped membranes. The red bamboo-like tissues and geometrically shaped membranes are both derived from tube-shaped structures. Our findings provide strong evidence that tube-shaped structures are stem cell niches. They are composed of only one type of lineage-specific stem cells, soluble, and insoluble matrix materials. In addition, our data provide evidences that the newly produced cellular progenitors, although contain specific nuclear materials, do not have unclears. Thus, the mechanism of stem cell self-renewal is not by mitotic division.

## Methods

### Animals and materials

Male Balb/C and C57BL6 mice at 8 to 16 weeks of age were purchased from Vital River Laboratories (Beijing). The mice received food and water *ad libitum*. The animal committee of Khasar Medical Technology approved the research protocol “stem cells in mouse blood”. All animal procedures followed the guidelines of the Animal Regulatory Office and relevant national and international guidelines. The anti-c-kit antibody was purchased from Santa Cruz Biotechnology (Santa Cruz, CA, USA). Donkey anti-mouse IgG was purchased from Invitrogen (Carlsbad, CA, USA).

### Mouse blood collection

Blood was collected from 5 of 8- to 10-week-old mice of each strain via cardiac puncture under sterile conditions after CO^2^ euthanasia. Each syringe was prefilled with 100 μl heparin (0.5%) to prevent coagulation. Blood was placed on ice for ∽5 min, pooled (∽5 ml), transferred to a 50-ml centrifuge tube, immediately diluted with 1x phosphate-buffered saline (PBS) at a ratio of 1:5 (blood:PBS), and then centrifuged at 200xg for 5 min. After removing the supernatant, the cellular portion was diluted with PBS at a 1:1 ratio, and then 15% paraformaldehyde was slowly added to a final concentration of 4% paraformaldehyde. Blood cells were fixed at 4°C overnight. The next day, the cells were centrifuged at 200xg for 5 min, washed 2 times with PBS and centrifuged at 100xg for 5 min and twice at 50xg for 5 min to remove free erythrocytes. Visible red clots were removed during washing. The cellular portions obtained after each centrifugation were pooled and examined. After a final centrifugation and removal of the supernatant, the cellular portion was fixed again with 4% paraformaldehyde at a ratio of 1:5 (original blood volume:fixative volume) and dropped (200 μm/drop from ∽5 cm above) onto gelatine-coated histology slides. At least 20 slides from each preparation for each strain of mice were stained and examined.

### Immunohistochemical staining

For immunohistochemical staining, the slides were blocked for 1 hr and incubated with an anti-c-kit antibody overnight at 4°C. After washing, the slides were incubated with a horseradish peroxidase-conjugated secondary antibody for 1 hr, stained with DAB, and then counterstained with haematoxylin before final mounting. The cells were visualized by conventional fluorescence microscopy (Leica, Wetzlar, Germany) and photographed using a digital camera (Leica, DFC500). The control tissue was treated with nonspecific antiserum of the same species.

### Haematoxylin and eosin (H&E) staining

Cellular portions from each mouse strain preparation were dropped onto at least 20 slides. The slides were dried overnight, washed 3 times with PBS, and stained with H&E. The stained slides were visualized by standard fluorescence microscopy (Leica) and photographed with the use of a digital camera (Leica, DFC500).

## Results

### Tube-shaped structures can be as long as 4 mm and various colours to H&E stains

The tube-shaped structures could be as long as 4 mm (Fig. 1A, 1C) that are embedded in the connective tissues (Fig. 1C). The width of these tube-shaped structures varied from 10 μm (Fig. 1D) to 50 μm (Fig. 1B), which may be due to the size and their maturation status.

**Figure 1.**
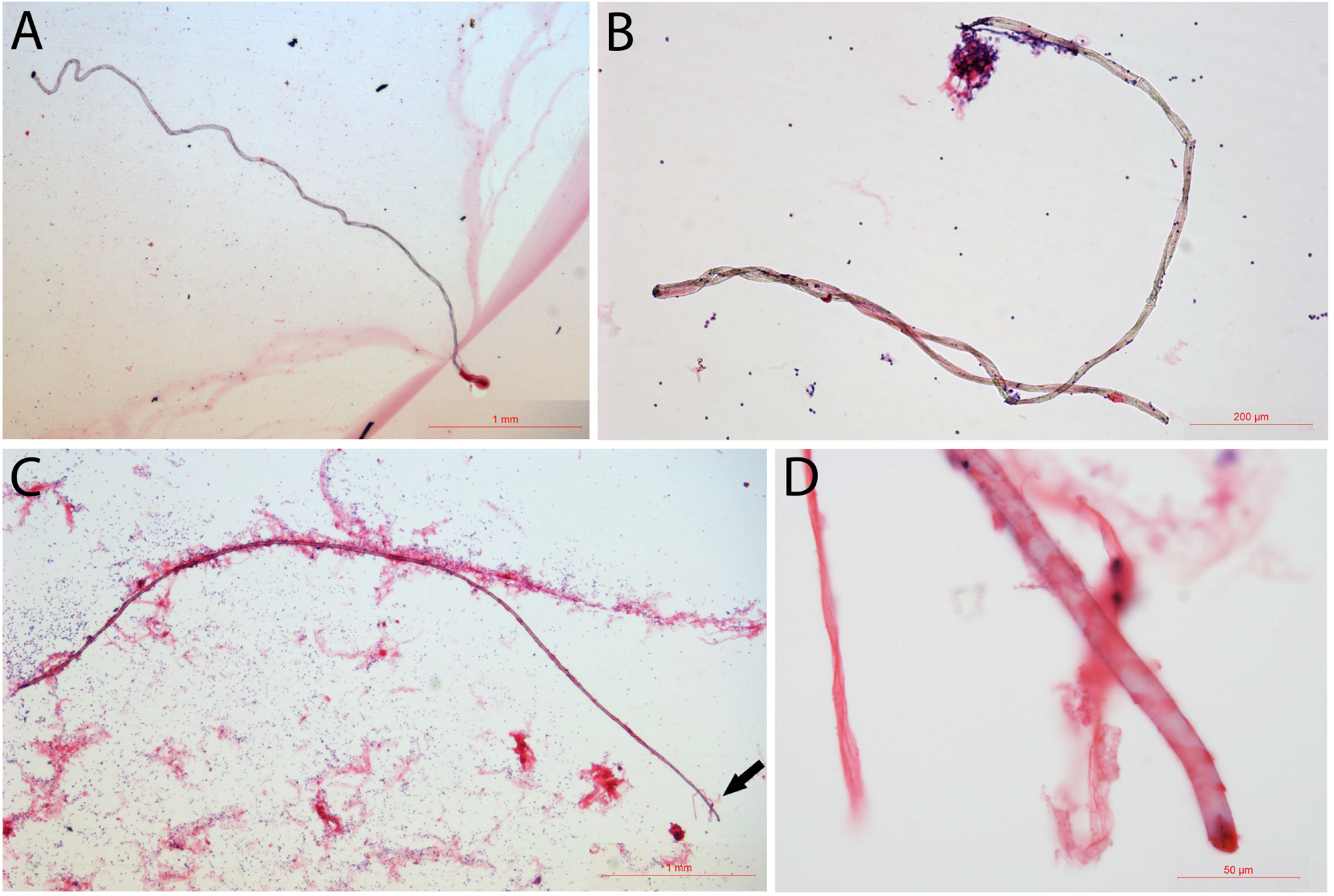
Tube-shaped structures in mouse blood. Tube-shaped structures can be as long as 4 mm (A - C). One of the tube-shaped structures was embedded in the connective tissues (C). The width of the tube-shaped structure was approximately 10 to 50 μm. Panel D shows the amplified image of the tip of the structure in panel C (arrow in C).

Haematoxylin and eosin (H&E) staining was performed to further examine the colour differences of these tube-shaped structures. Besides the light-purple colour, copper colour, and dark blue colours that were described in our previous publications ^19, 22^, we identified more colours stains on these tube-shaped structures. Figure 2 reveals several tube-shaped structures sorted by the H&E staining: dark-grey (Fig. 2A), dark blue (Fig. 2B), no staining (Fig. 2C), and red colour (Fig. 2D). Thin filaments can be observed inside the dark-grey coloured (arrow in Fig. 2A), suggesting that it had thin filament-like tissues inside. The dark-blue one (Fig. 2B) had a spiral opening along the tube, suggesting that it has released its inclusions. Figure 2C is a no-staining tube-shaped structure that held numerous tiny particles inside. A short horizontal line could be seen in the middle (arrow in Fig. 2C) of it. Also, thin filament-like tissues were observed at the end of it. The red-coloured one (Fig. 2D) that was more than 1 mm in length had many crossing thin lines (arrows in Fig. 2D). These crossing lines had the similar distances to each other, suggesting fragmented tissues inside. None of these tube-shaped structures were related to blood vessels. We believe that although H&E staining had limitations in identifying these tube-shaped structures, it is currently the only available method to sort these tube-shaped structures.

**Figure 2.**
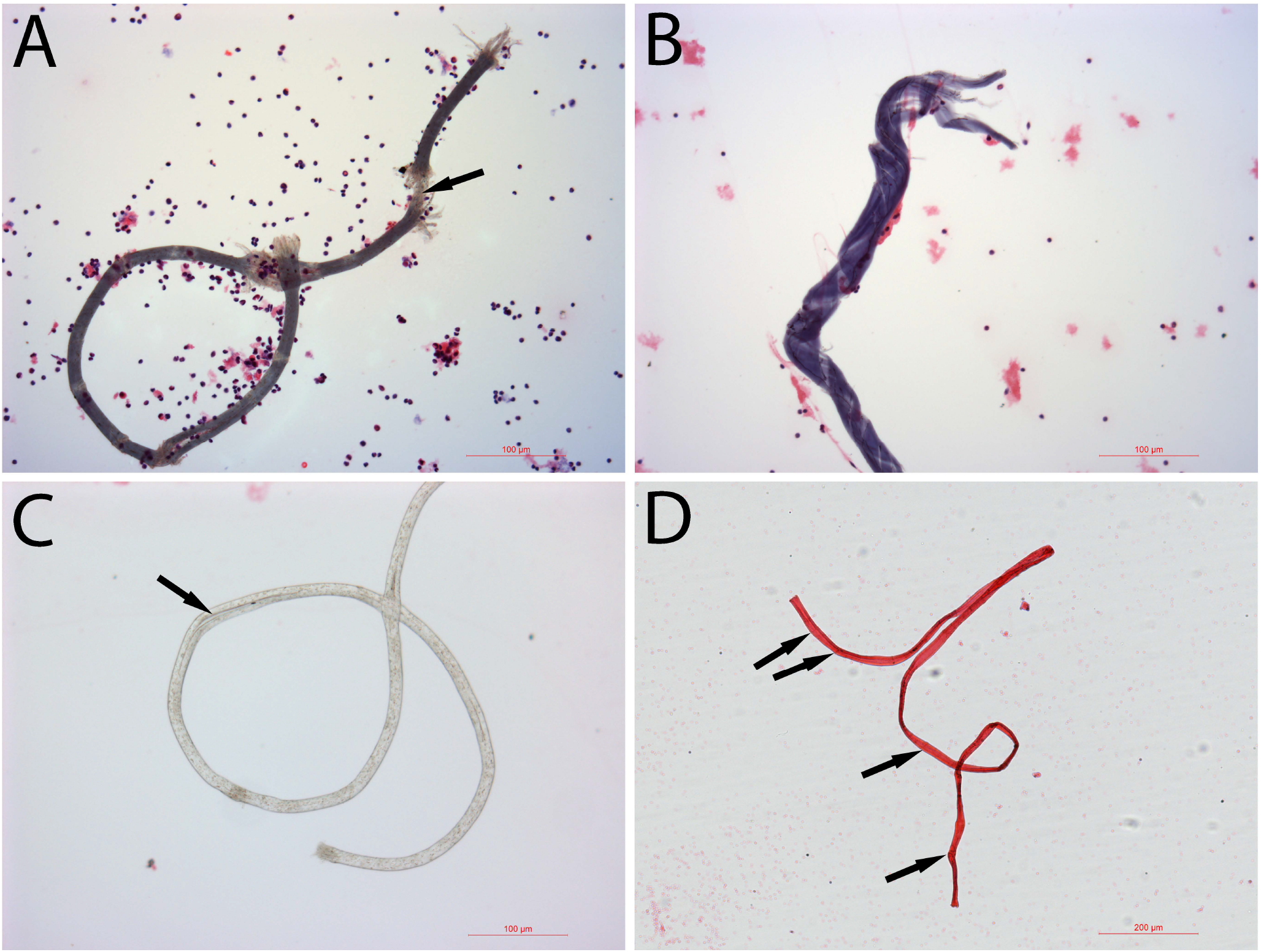
H&E staining of the tube-shaped structures. H&E staining showed that some tube-shaped structures were dark grey (A), dark blue (B), not stain (C), and crimson (D) Bars in A to C = 100 μm; bars in D = 200 μm.

### Red bamboo-like inclusion (RBLIs)

A tissue that was approximately 1.4 mm in length, stained red, and had bamboo-like joint sections were identified (Fig. 3A). Higher-magnification images on both ends showed that was connected by the sections to each other with an overlapping knot (Fig. 3B, Fig. 3C). Additionally, higher-magnification images revealed many dark particles on the red bamboo-like tissue (arrows in Fig. 3B, Fig. 3C). These dark particles had the same colour stains of the adjacent nucleus, suggesting that they were nuclear materials. Then, many red membranes were identified. Some of them were still in the folded shape (arrows in Fig. 3D). Red membranes were approximately 30–40 μm in diameter, which was similar to the length of each red segment in the bamboo-like tissue, suggesting that these red membranes were unfolded after splitting from the bamboo-like tissues. Additionally, a line of split red membrane progenitors was located on the top of a line of jelly-like tissue (arrows in Fig. 3E), further suggesting that these red membrane progenitors were from the bamboo-like tissues that had the jelly-like transparent outer layers.

**Figure 3.**
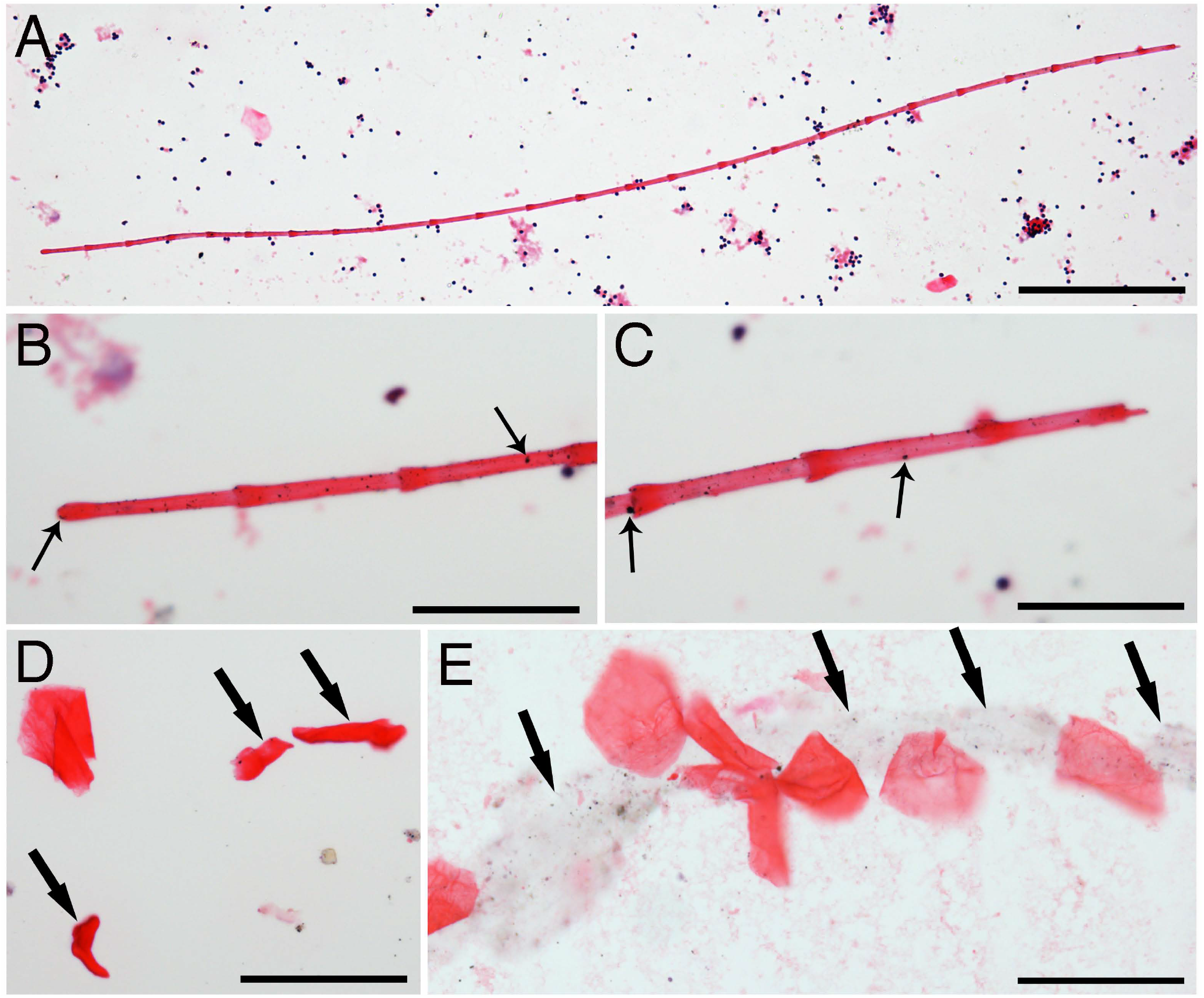
Identification of RBLIs. One red bamboo-like structure was approximately 1.4 mm in length, stained red and had joint sections (A). Higher magnification images of its both ends showed the arrangement of the sections, and many dark particles on them (arrows in B, C). Some red membrane stem cell progenitors were still folded (arrows in D). Additionally, a line of jelly-like tissue (arrows in E) was found beneath the extended red membrane progenitors. Bar in A = 200 μm; bars in B-E = 50 μm.

Red thin membrane-like structures can be abundant in the blood (Fig. 4A). Although many nuclei can be seen together with these red thin membranes (Fig. 4A), none of the nucleus belongs to the thin membranes. Figure 4B showed that, in a group of purified red thin membrane structures, no nucleus appeared in any of them. We have noticed that even though the red thin membranes are abundant in the blood, they were not been described or examined in haematology studies. The reasons for their not being seen could be due to their slippery or non-stick peculiarity on tissue slides, and due to their light-weight, which makes them stay mostly in the serum and red blood cell levels during gravity blood separation. Also, we believe that RBLIs differentiate after being released from the tube-shaped structures. Supplementary Figure 1 shows an RBLI that is about 1 mm in length and has almost no stain to H&E. High magnified images (SF1B, SF1C) revealed that faint pink colour stained only on the overlapped areas. Not staining to H&E has been described in thin filaments and fusiform-shaped structure-derived progenitors. This data further support that the inclusions and pre-stem cell progenitors continue their development after being released.

**Figure 4.**
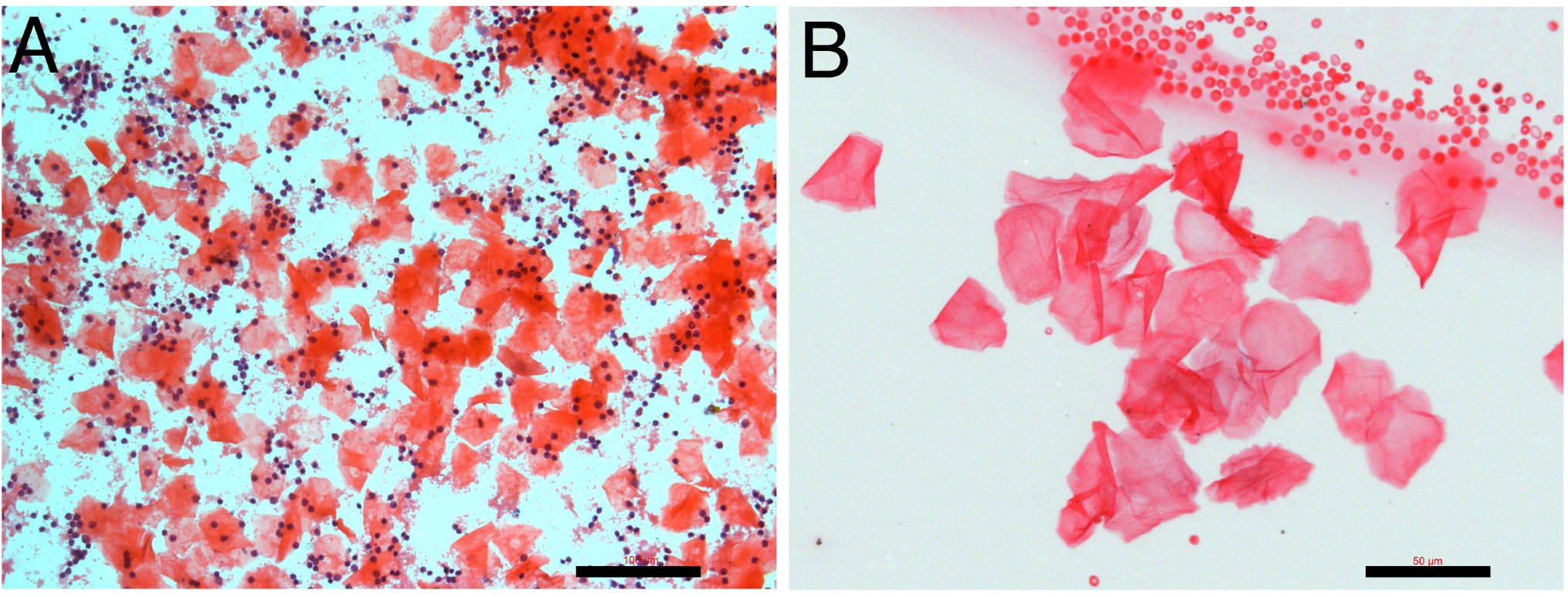
Red membrane-like progenitors are abundant in the blood. Many red membranes can be observed together with numerous monocytes (A). Grouped red membranes without nucleus were isolated (B) together with small red blood cells adjacent.

### Nuclear materials in the red membranes

We then asked whether these red thin membranes are pre-stem cell progenitors if they do not have a nucleus. Higher-magnification images revealed that although none of them had a nucleus, every one of them contained tiny dark particles (arrows in Fig. 5A-5D) or DNA fragments. In the early stages, only a few small DNA particles (arrow in Fig. 5A, 5B) were seen. The more developed red membranes (Fig. 5C, 5D) showed some red colour contrasts that were similar to the grid patterns. Many DNA particles that aggregated in each grid were observed in each membrane (arrows in Fig. 5D). Red thin membrane-like cells were observed (Fig. 5E). This cell had a regular-sized nucleus, a small-sized nucleus and aggregated DNA particles (arrows in Fig. 5E). In addition, numerous DNA fragments could be seen in this red membrane-like cells. We believe that these DNA fragments guide and determine nucleus and cell lineage formation. Nucleus formation from DNA fragments has been described in our previous publications ^20^. Although we still do not know the mechanism of nucleus formation, we believe that DNA fragment polymerization and fusion may occur in this process.

**Figure 5.**
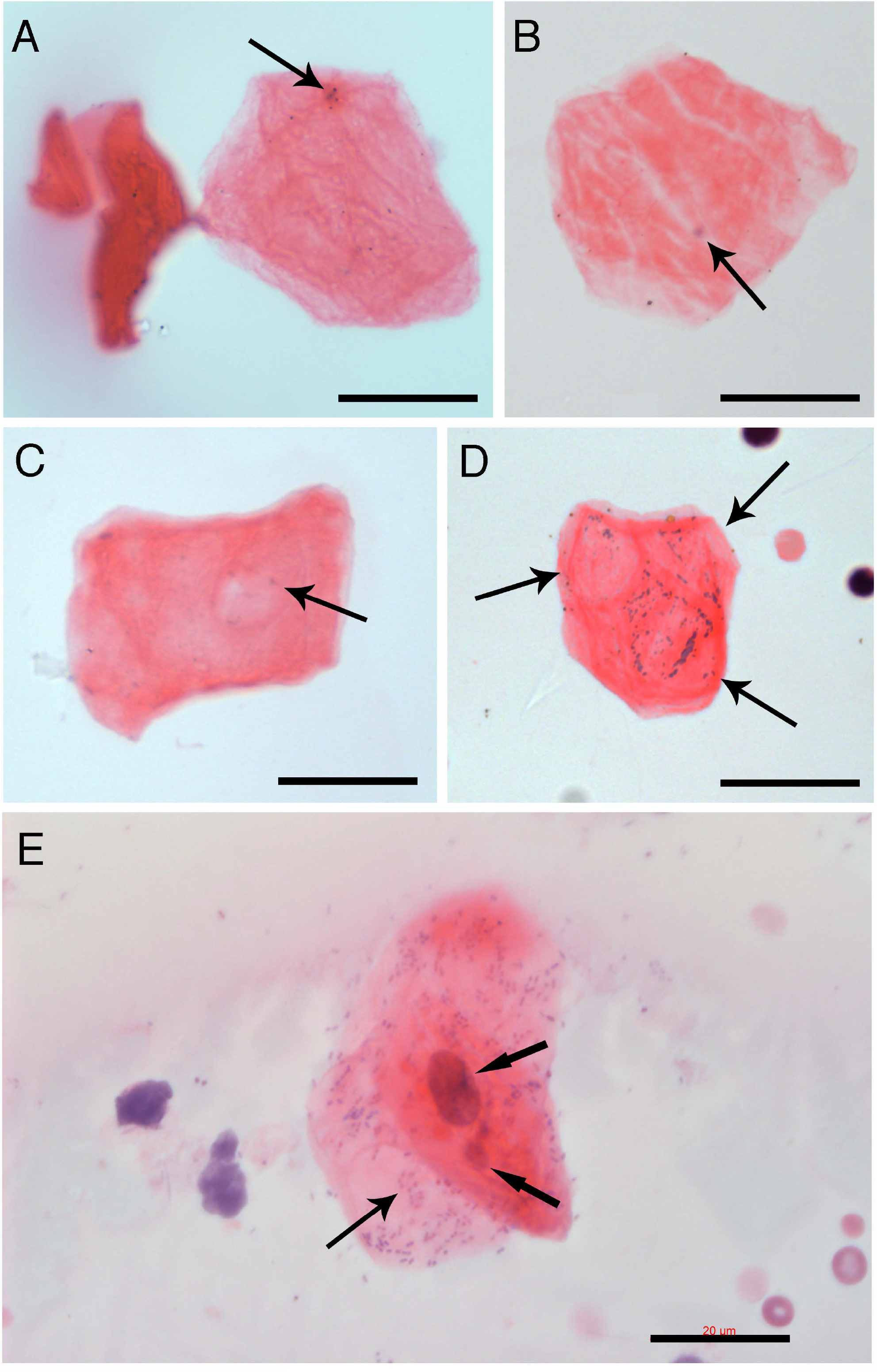
RBLI-derived progenitors. Higher magnified images revealed that every red membrane-like progenitors contained tiny, dark DNA particles (arrows in A-B). Red colour contrasts (arrow in C) appear on the membrane, which were similar to the grid patterns. Many DNA particles aggregated separately in three grid patterns of one red membrane (arrows in D). Red thin membrane-like cell had a regular-sized nucleus, a small-sized nucleus, and numerous aggregated DNA particles (arrows in E). Bars = 20 μm.

### Another tube-shaped niche releases geometrically shaped membrane inclusions

To support our evidence that the tube-shaped structures are lineage-predetermined stem cell niches, we present another tube-shaped structure that releases geometrically shaped membrane inclusions. Figure 6A shows a tube-shaped structure that was more than 2 mm in length, located parallel to a line of non-staining geometric-shaped inclusions. The morphology of these geometrically shaped inclusions was not like those we described in our previous publications or above. They did not stain with H&E, had different shapes, such as diamond, square, triangle, or irregular shapes, and ranged in size from 30 × 30 μm to 200 × 60 μm. These geometrically shaped inclusions were arranged in lines closely adjacent to the tube-shaped structure, which suggests that they are released from the tube-shaped tissue. They attached well to the tissue slides, suggesting that they are thicker and heavier than the red membranes described above. Higher-magnification images revealed that these geometrically shaped inclusions had many tiny cracks on their surfaces (Fig. 6B, 6C).

**Figure 6.**
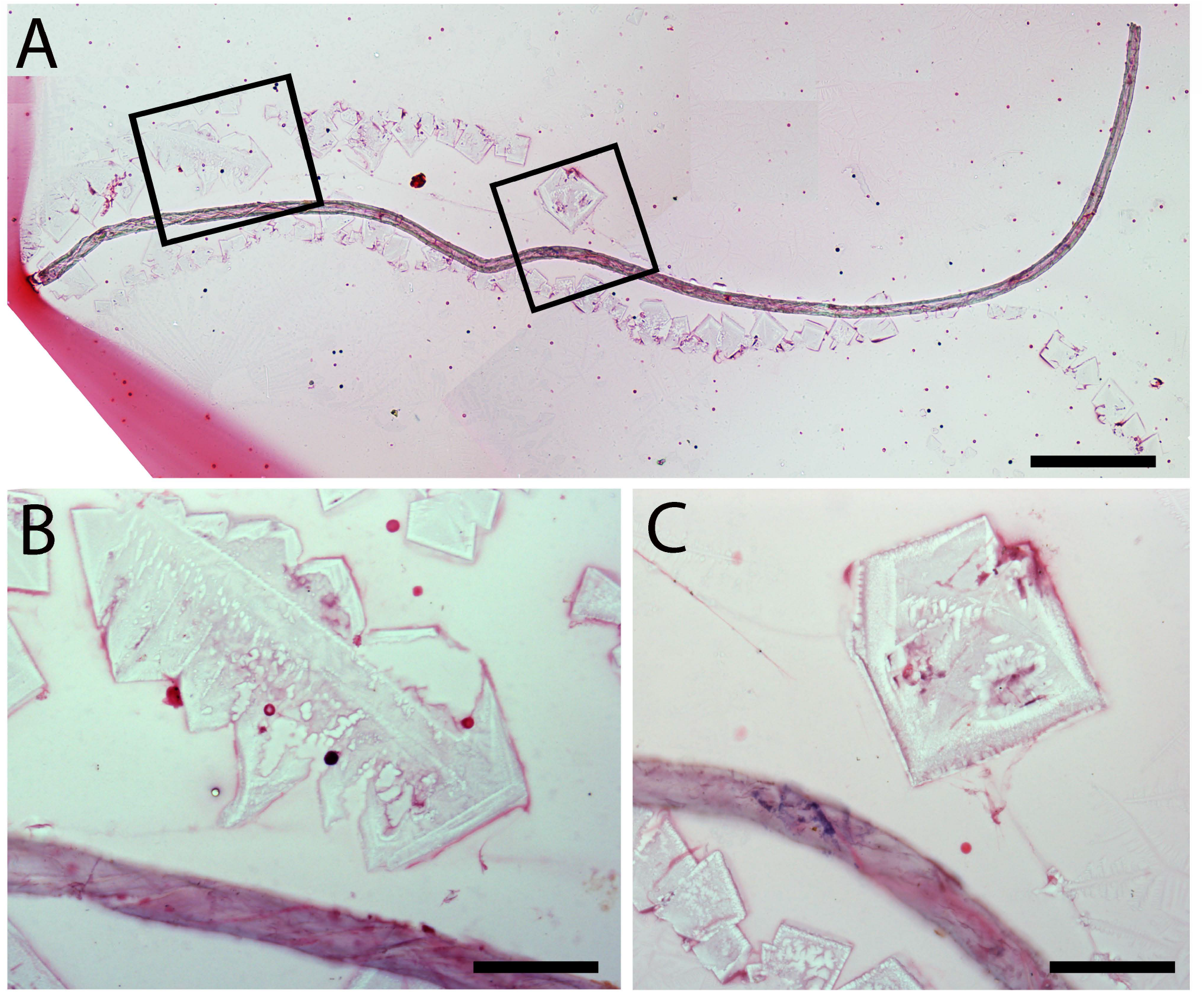
Geometric-shaped membrane inclusions. A tube-shaped structure that was more than 2 mm in length, located parallel to a line of non-staining geometric-shaped inclusions (A). Higher-magnification images revealed that these geometrically shaped inclusions had many tiny cracks on their surfaces (B, C). Bar in A = 200 μm; bars in B and C = 50 μm.

### Small pre-stem cell progenitors developed inside geometrically shaped inclusions

We then identified similar geometrically shaped inclusions that stained slightly blue and were square- (Fig. 7A), diamond- (Fig. 7B, 7C), or triangle-shaped (Fig. 7D-7F). They sized about 400 × 500 μm, which were much larger than the newly released ones described above, which suggest that these membranes can become larger after being released, although smaller-sized membranes can still be seen (Fig. 7A, 7C). On the larger-sized membranes, some areas were not stained, but most areas had numerous small dot-shaped progenitors that had weak blue staining.

**Figure 7.**
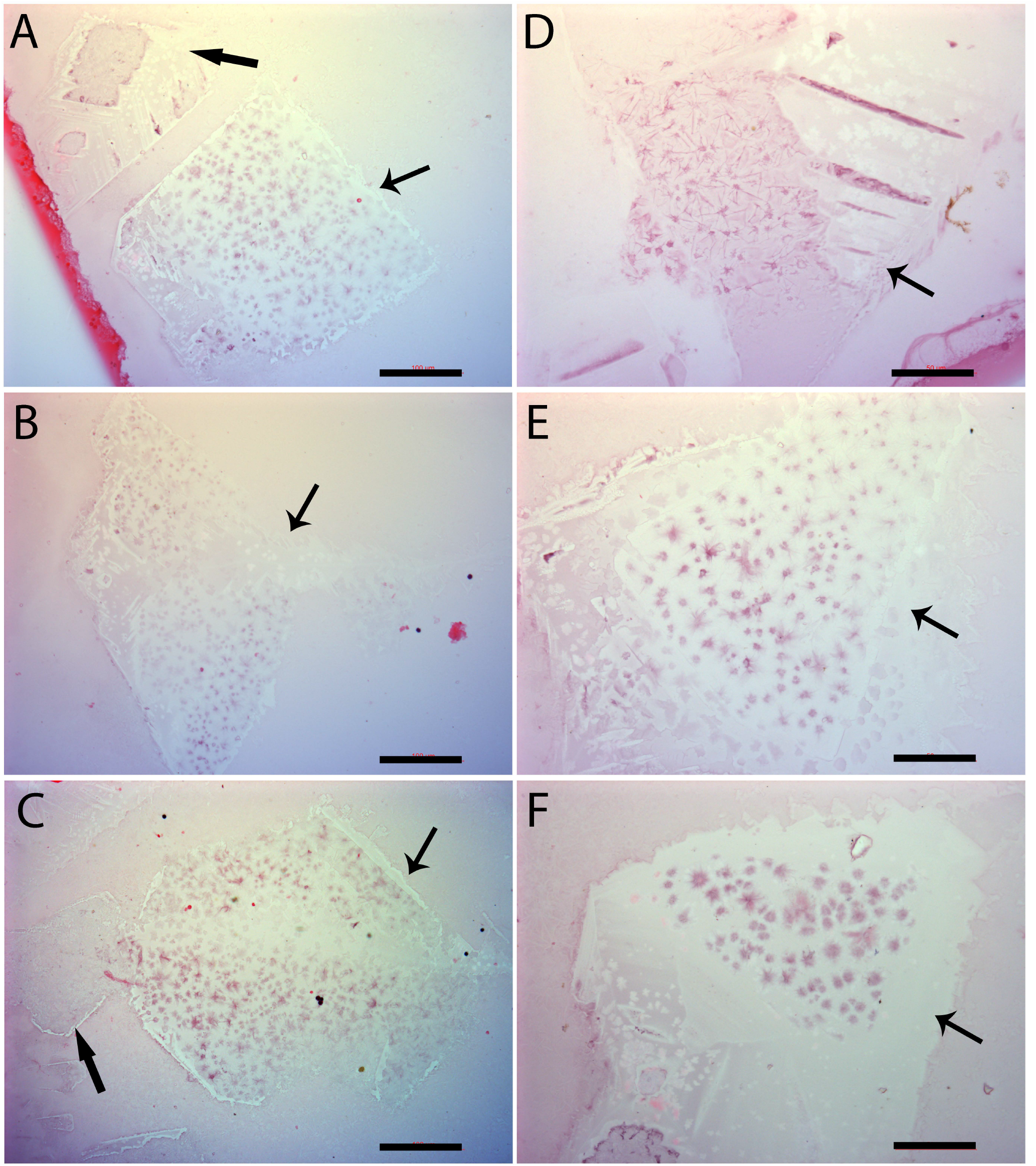
Geometric-shaped inclusion-derived progenitors. H&E stains revealed morphologically similar geometrically shaped inclusions. They were square- (A, C), diamond- (B), or triangle-shaped (D - F). Some areas on their surfaces were still not stained (arrows in A, B, C), but most areas were stained slightly blue, with staining more concentrated on the small dots. Small geometric-shaped inclusions had almost no stains (thick arrows in A, C). Bars in A-C = 100 μm; bars in D - F = 50 μm.

Higher-magnification images revealed that small dot-shaped cellular material that sized about 4 to 7 μm in diameter were formed inside these geometrically shaped membranes (Fig. 8A - 8D). Tiny cracking lines around each small dot-shaped cellular structure were observed (arrows in Fig. 8A, 8B), which suggest that the membrane breaks to release these small dot-shaped cellular structures. The released small progenitors stained blue to H&E and had a size of approximately 5 μm (Fig. 8C, 8D). We hypothesize that after being disconnected from the membrane, these dot-shaped cellular structures can further undergo differentiation, including nucleus formation and specific cellular membrane formation, to become nucleated cells. Thus, dozens to hundreds of small progenitors can be produced in one membrane. We also noticed that most small blue progenitors appeared earlier in the center of the membrane than in the peripheral areas.

**Figure 8.**
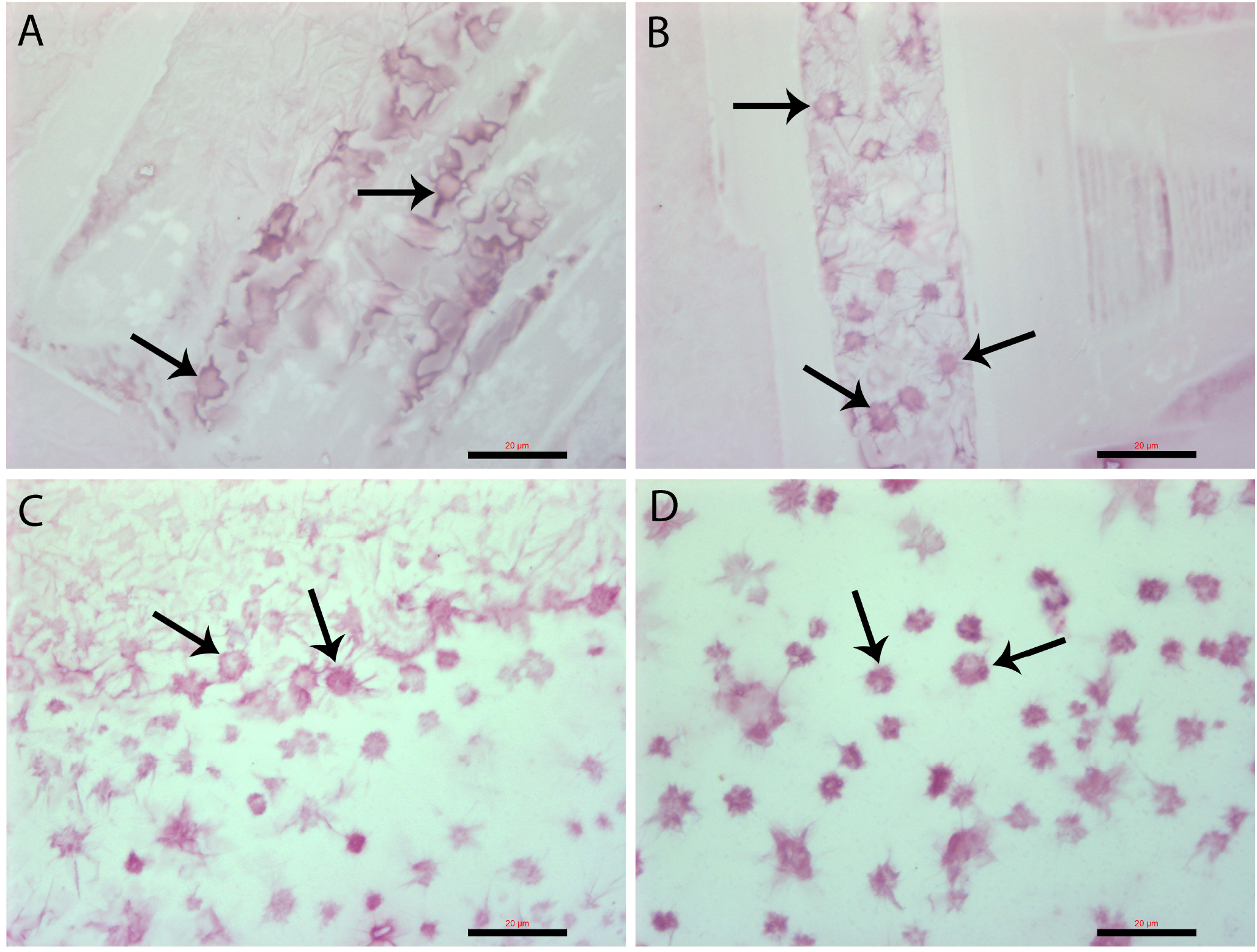
Geometric-shaped inclusion-derived progenitors. Higher-magnification images revealed that in the early stages, the surfaces of the inclusions were tiny circles surrounded by small lines (arrows in A, B). The tiny circles cracked to release individual small progenitors (arrows in C, D). These small dot-shaped progenitors were about 3 to 7 μm in diameter. Bars = 20 μm.

### The geometrically shaped membrane-derived small blue progenitors are C-kit positive

C-kit has been used as a stem cell marker to study or sort stem cells. Therefore, the expression of C-kit on the geometrically shaped inclusions was investigated. The images (Fig. 9A, 9B) showed that c-kit was specifically expressed on the small progenitors located in the center areas of the inclusions, which also supports the idea that the small progenitors in the center were more mature or more differentiated.

**Figure 9.**
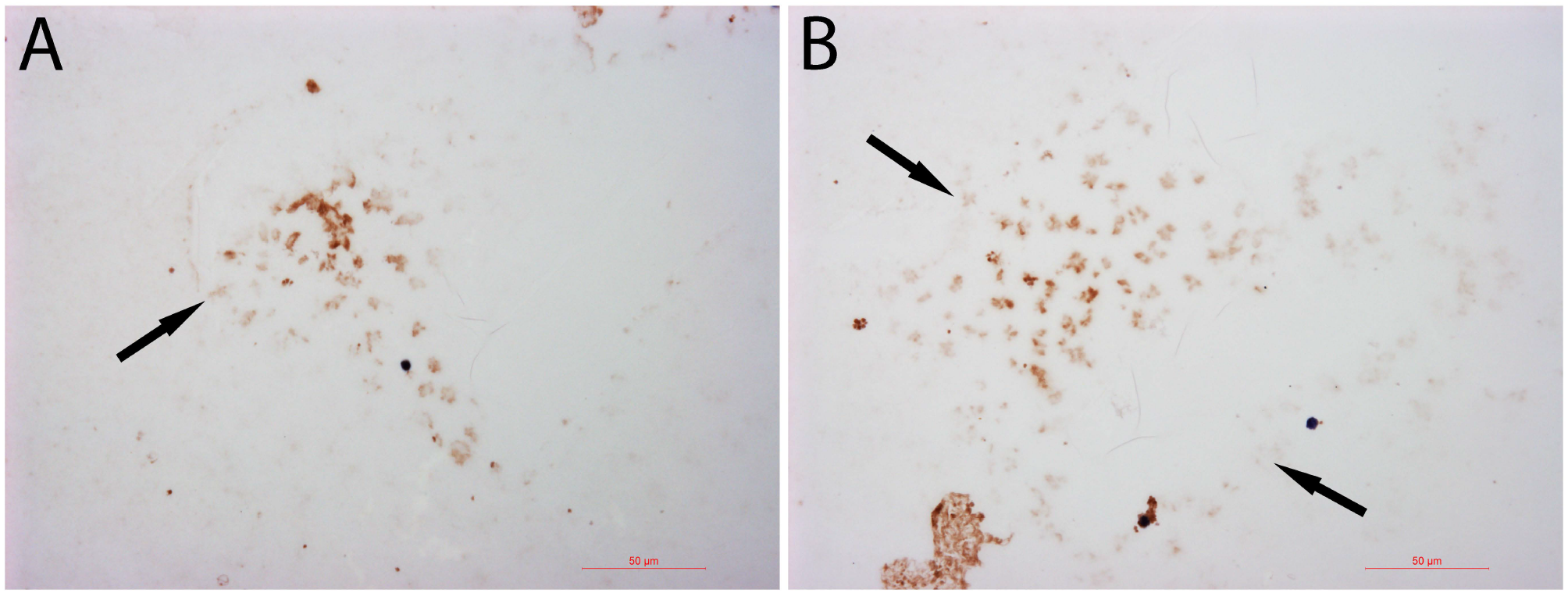
Geometric-shaped inclusion-derived progenitors were c-kid positive. C-kit was specifically expressed on only the small progenitors located in the centre of the inclusion (thick arrows in A, B). Bars = 50 μm.

## Discussion

In our previous studies ^19 22^, we reported three H&E-stain distinct tube-shaped structures, a light purple colour, a copper colour, and a dark-stained colour, and their inclusions, including progenitor-releasing filaments, bud-shaped, and dark fragment structures. Now, we provide evidence of another two types of tube-shaped structures and the released inclusions, RBLIs, and geometric-membrane-shaped inclusions. The RBLIs released red membrane progenitors, and the geometrical-shaped membranes produced small c-kit-positive progenitors. Our data provide strong evidence that the tube-shaped structures are lineage-restricted stem cell niches.

Each tube-shaped structure releases morphologically distinct inclusions, which suggests that their specificities have been predetermined. The tightly enclosed long tube-like structure suggests that any cells or factors inside the lumen are completely isolated from the outside environment. In the previous report ^19^, we found that some nucleated cells were located inside the tube-shaped structures. Although we could not identify any nucleated cells located with RBLIs or geometric-shaped membranes in the current study, we still believe that the lineage-restricted nucleated cells are the original resources that build and determine the lineage of the tube-shaped structures. One nucleus of these cells could release hundreds of DNA fragments that become genetic materials to initiate the formation of many restricted progenitors. Thus, one cell lineage with numerous DNA fragments, and specific cellular molecules that reside inside the niche participate in niche assembly and specific stem cell progenitor development. Other types of cells are not present or involved in the development of these niches. We believe that these tube-shaped niches are built in specific cellular areas, from where they can migrate to the circulation to release their contents. Our belief is based on a previous report that erythrocyte sacs ^21^, the inclusions of red blood cells, are found only in the blood but not in the bone marrow.

Unlike the tube-shaped niches, each restricted inclusion showed not only a colour difference but also fundamental morphological differences, such as the thin filaments ^19^, bud shaped ^19^, erythrocyte sacs ^21^, dark fragments ^22^, plus the currently described red bamboo-like and geometric membrane-shaped. Thus, each lineage was sorted out by its unique colour and distinct morphology. Many of them are difficult to detect because they do not stain with H&E, let alone any specific markers. Except for jelly-like materials, we still do not know what the outer layers of the inclusions are composed of. Our data show that each of these inclusions can produce dozens to hundreds of progenitors in a synchronized pattern. Some progenitors reached maturation before being released from the inclusion, such as red blood cells ^21^; some continued to differentiate even after being released from the inclusions, although the mechanism of nucleus formation in these enucleated progenitors is still unknown.

A recent report indicated that lineage-restricted segregation occurred during early embryonic development ^17^. In later embryonic development or postnatal life, stem cells are only unipotent. Our data showing that each tube-shaped niche produces only one type of stem cell progenitor strongly supports this report. We believe that lineage segregation occurs during early embryonic development, so only lineage-restricted cells exist in postnatal life. The life-long production of unipotent progenitors in postnatal life occurs in tube-shaped niches that are formed by specific unipotent cells containing numerous DNA fragments. This renewal process occurs via a recurrent type. New progenitors are formed in their niches via the combination of lineage-restricted DNA fragments, cellular proteins, and other relevant factors. Stem cells were renewed inside tightly sealed tube-shaped structures and morphological distinct inclusion, which suggests that there is double protection for fragile and/or tiny stem cell progenitors during their development. The further differentiation of these progenitors occurs in the circulation, where many factors, including cellular proteins, nucleic acids, regulatory factors, and even specific cellular membranes, participate in the formation of nucleated cells.

We termed the newly developed enucleated cells “progenitors” because they are unipotent, and are upstream of unipotent nucleated cells. Our current data indicate that restricted stem cell progenitors are more primitive than nucleated stem cells because they do not have nuclei and can differentiate into nucleated cells or unipotent stem cells. The differentiation of progenitors occurs after they are freely released in the blood or specific destinations. Since these newly produced progenitors contain only DNA fragments or do not have nuclei, they cannot replicate unless they become nucleated stem cells or fully differentiate. We believe that when nucleus formation is completed, mitotic division begins.

Our data showed that c-kit, which has been extensively used for stem cell sorting, was expressed on newly produced small blue enucleated progenitors. Thus, we assume that using antibodies in cell sorting, the method used by many researchers could sort out enucleated progenitors and not nucleated cells. On the other hand, many progenitors, such as red membrane-like or early-stage small blue progenitors, do not express stem cell markers on their surfaces and cannot be sorted out by any known stem cell markers. Thus, we can conclude that even though cell sorting has been used extensively to speed up data acquisition in many investigations, it has obvious limitations in stem cell research. Most newly produced enucleated progenitors, regardless of lineage, cannot be selected by cell sorting unless they have reached certain maturation stages.

Because most of the released inclusions and the majority of small progenitors are either small in size or light in weight, they are located in the blood at the similar density to platelets or serum before their full differentiation. Thus, they could be mistakenly classified as platelets, and their regenerative functions could also be incorrectly credited to platelets.

## Acknowledgements

This work was supported by a grant from the Department of Technology, Inner Mongolia, China.

## Competing interest

None of the authors have any competing interests for the work in this manuscript.

## Author Contributions

W. Kong: Manuscript writing, Project directing, Experimental designs.

X. Han: Performing experiments, Data collection.

H. Wang: Performing experiments, Data collection.

X. Zhu: Performing experiments, Data collection.

## Notes

### Competing Interest Statement

The authors have declared no competing interest.

