## Supplementary material for "Development of membrane-like pre-stem cells after released from tube-shaped niches": Supplemetary Figure

Supplementary Figure

Supplementary Figure 1. Bamboo-like structures develop after their releases. One bamboo-like structure was approximately 1 mm in length was identified (A). Higher magnification images of its both ends showed pink stains only on the overlapped areas but not whole sections (B, C). Bar in A = 200 μm; bars in B-C = 20 μm.

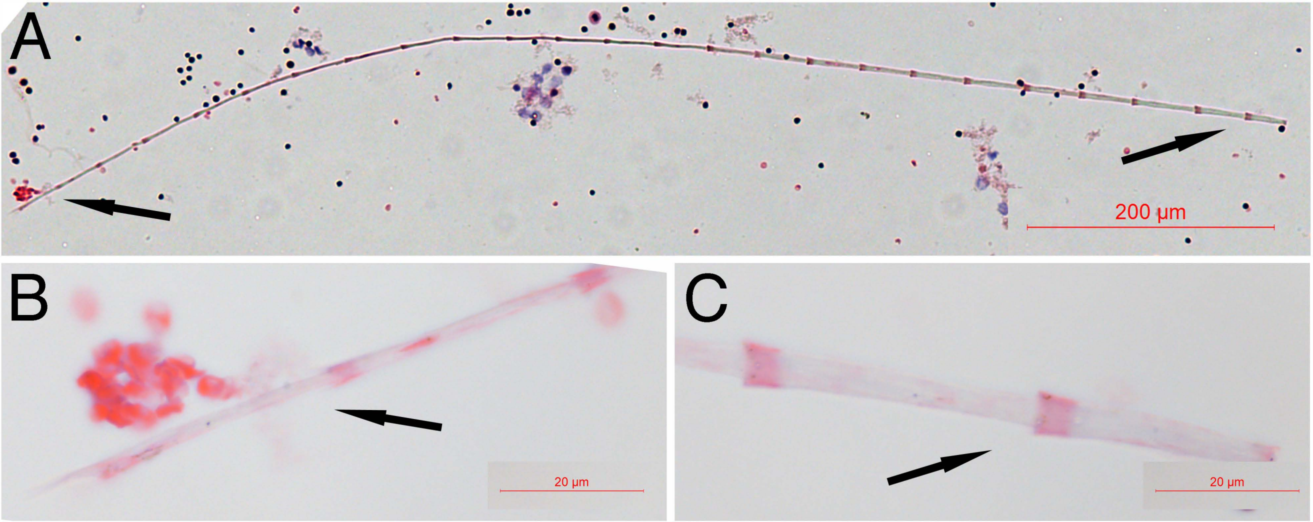


Supplementary Figure 1
